# Projected Gradient Descent for Hallucination-Based Protein Binder Design

**DOI:** 10.64898/2026.10.22.760942

**Authors:** Magnus Ross, David Miller, Jude Wells

## Abstract

Gradient-based hallucination is a popular approach to protein binder design that backpropagates a loss defined on a structure predictor’s output to guide optimisation in a continuous relaxation of sequence space, yet the choice of optimiser for this task has received little systematic evaluation. In this work we evaluate simplex-projected gradient descent (PGD) as an alternative to the commonly used gradient descent on softmax-reparameterised sequence logits. By updating amino-acid probabilities directly, PGD avoids softmax-induced gradient attenuation and allows amino acids suppressed early in optimisation to re-enter the design. Using an ESMFold2-based pipeline on ten targets, we find that PGD minimises the design loss in fewer iterations and produces designs with higher predicted interface confidence compared to its softmax counterpart. Simulating selection in real campaigns by considering the upper tail of the ipSAE distribution across a pool of 1,024 designs per target, we show that PGD significantly improves on six of the ten targets, with no significant degradation on any, and produces more structurally diverse designs on every target. Additionally, we find that pairing shorter PGD runs with candidate screening raises the total number of in silico hits per GPU-hour by 29% relative to default-length trajectories. These results support PGD as a simple and effective alternative for hallucination-based binder design. Code and design data are publicly available.

**Code** github.com/magnusross/pgd-hallucination

**Data** doi.org/10.5522/04/34493751

## 1 Introduction

Protein binder design aims to create proteins that bind a target protein of interest with high affinity and specificity, with uses spanning therapeutics, diagnostics, and synthetic biology. Traditionally this meant experimentally screening thousands of candidates with expected hit rates under 1% (Cao et al., 2022). Hit rates improved markedly with the arrival of reliable deep-learning-based structure prediction models. The confidence scores these models assign to a predicted complex are predictive of experimental binding (Bennett et al., 2023; Claude Science & Shanehsazzadeh, 2026; Overath et al., 2025), so candidates can be ranked in silico and only the top-ranked designs sent for experimental validation; we review the evidence for these metrics as predictors of binding in appendix G. Hits can now be expected from around ten tested sequences, with a recent campaign reporting confirmed binding for 27% of tested designs across 15 targets (Claude Science & Shanehsazzadeh, 2026). Nevertheless, the computational efficiency with which these designs are generated remains a concern. Typically, the number of in silico candidates generated is many times larger than the number that go on to be tested in the lab, with recent campaigns requiring hundreds of H100 GPU-hours per validated hit (Candido et al., 2026; Claude Science & Shanehsazzadeh, 2026). Computational binder design can then be framed as the task of producing, as cheaply as possible, sequences that score well under these predictive in silico metrics. The space of possible binder sequences is far too large to search exhaustively, but since the scoring metrics are built from a differentiable structure predictor, high-scoring sequences can be found by inverting the structure predictor and performing gradient descent in a continuous relaxation of sequence space, in which each position in the designed binder carries a distribution over the 20 standard amino acids (Frank et al., 2024). This is known as *hallucination-based binder design*, and is the approach we focus on in this work. The soft sequence being optimised therefore lives in a space defined as the product of probability simplices, one simplex per sequence position, and the search is a constrained optimisation in that space whose final iterate will be discretised into a sequence.

A number of works have proposed advances in hallucination-based binder design, including the addition of filtering (Pacesa et al., 2025), new loss terms (Nori et al., 2025), and more efficient software implementations (Boyd et al., 2025). One area that we believe to be of underappreciated importance is the optimisation algorithm itself. Whilst a number of approaches have been used in the literature, the optimiser in each new work is subtly different, and these optimisers are introduced alongside a slew of other changes to the methodology, making it difficult to isolate the effect of the optimiser. Innovations in optimisation algorithms for neural networks have led to large performance improvements (Kingma & Ba, 2015; Liu et al., 2025), but the structure of the problem in the binder design context is distinct. The space is much lower-dimensional and is typically constrained to the simplex (Boyd et al., 2025; Frank et al., 2024). Taking gradient steps while satisfying the simplex constraint is the problem that gives rise to the different optimisers we consider in this work.

We use the design pipeline proposed by Candido et al. (2026), as the ESMFold2 family of models achieves state-of-the-art performance in structure prediction and attained high experimental hit rates in the accompanying binder design campaign. We replace their softmax-reparameterised gradient descent, which we call Logit GD, with simplex projected gradient descent (PGD) (Boyd et al., 2025; Calamai & Moré, 1987), which projects the iterates back onto the simplex only after steps that violate the simplex constraints (fig. 1). The softmax reparameterisation, used by many approaches (Frank et al., 2024; Pacesa et al., 2025), satisfies the constraint without the need for projection, but the logit gradient for an amino acid shrinks as its probability approaches zero (Linder & Seelig, 2021), so choices that lose probability early are, in practice, never recovered (Mei et al., 2020). PGD instead updates the probabilities directly, avoiding this vanishing-gradient problem and ensuring that low-probability amino-acid identities remain reversible (fig. 5c). Our paper makes the following contributions:

- We frame hallucination-based binder design as a constrained optimisation problem over a product of simplices, and show some of the limitations of the commonly used softmax reparameterisation, motivating simplex PGD as a drop-in alternative.
- We evaluate in silico simplex PGD against the softmax-reparameterised Logit GD of the ESMFold2 pipeline on ten targets with 1,024 designs per target and optimiser. We release all generated binder sequences as well as predicted structures and optimisation trajectories (Ross, 2026).
- We show that PGD gives significantly improved in silico binder confidence scores on six of ten targets, significantly lower confidence scores on none, and that these designs are more structurally diverse on all ten targets.
- We show that combining shorter PGD runs with candidate screening increases the pooled number of in silico hits per GPU-hour by 29% relative to longer PGD runs, across ten targets.
- We analyse the optimisation dynamics and provide evidence that PGD can explore sequence space more extensively than Logit GD.

**Figure 1:**
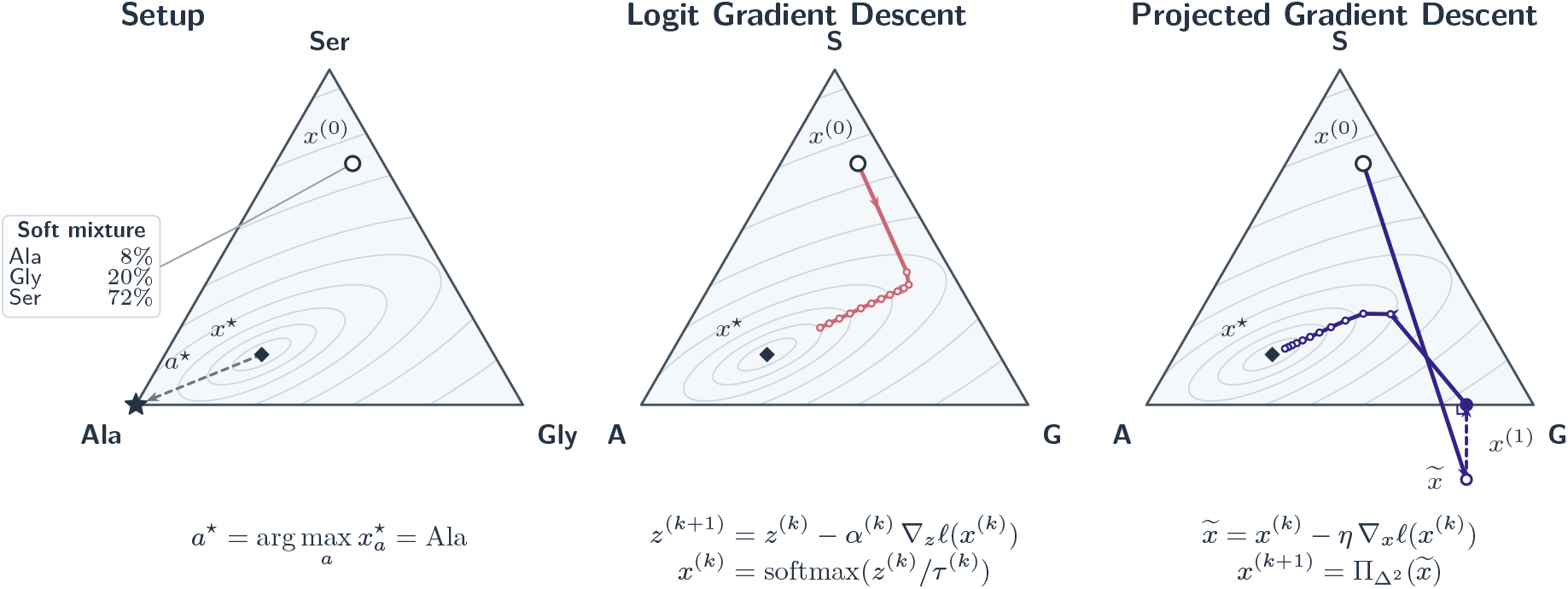
Optimisation geometry on the probability simplex. Hallucination-based binder design can be framed as gradient descent on the probability simplex, with points representing a soft mixture of amino acids, followed by discretisation of the converged iterate. For ease of visualisation, here we show the optimisation problem for a binder with a single position, and an alphabet of only three amino acids, implying *x* ∈ Δ^2^. The open circle is the initialisation *x*^(0)^, the diamond the global optimum *x*^*⋆*^, and the star the discretised residue *a*^*⋆*^ = arg max_*a*_ 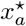. Logit GD updates logits *z* with step size *α*^(*k*)^ and maps them to the simplex by softmax at temperature *τ*^(*k*)^; PGD takes a fixed-size step *η* to 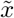, which may leave the simplex, and projects it back with the Euclidean projection 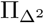. Approaches based on reparameterisation with softmax followed by gradient descent on the logits can suffer from vanishing gradients near simplex edges. Projected gradient descent in probability space avoids this issue.

## 2 Background

In this section we introduce the core problem of hallucination-based binder design, provide details on the construction of suitable objectives and discuss some algorithms for minimisation of these objectives.

### 2.1 Problem setup

A protein is a sequence *s* = (*s*_1_, …, *s*_*L*_) of length *L*, with each residue *s*_*i*_ drawn from the alphabet *A* of the 20 standard amino acids. Binder design is always posed relative to a fixed *target* 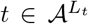, which is an input to the problem and is never optimised; we write 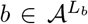 for the binder being designed. In the hallucination-based binder design framework we assume we have some loss function 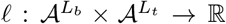 which determines the quality of a candidate binder *b* against the target *t*. To generate a single design, we wish to find a sequence *b*^*⋆*^ that minimises the loss for the given target, so

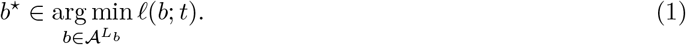

### 2.2 Relaxation

The optimisation problem in eq. (1) is discrete, and can be solved directly using discrete search algorithms such as MCMC in sequence space (Anishchenko et al., 2021). An alternative approach is to solve a continuous relaxation (Frank et al., 2024). Discrete protein sequences must be embedded in continuous space before deep-learning-based structure prediction models can be applied. This is achieved by encoding each residue in the sequence as a one-hot vector. Thus the sequence *s* is represented as a matrix *S*∈{ 0, 1} ^20*×L*^ whose columns are these one-hot encodings. Although superpositions of amino acids are not part of the training distribution, it is also possible to supply them at each position, so that the input to the models, representing the candidate binder sequence, is a matrix 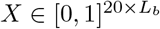 in which each column is a normalised probability vector that lies on the simplex Δ^19^ = {*x* ∈ R^20^ : *x* ⪰ 0, **1**^⊤^*x* = 1}. Following common usage in the protein design literature (Frank et al., 2024; Goverde et al., 2023; Norn et al., 2021) we refer to *X* as a position-specific scoring matrix (PSSM), though we note that it is a position-specific *probability* matrix. Frank et al. (2024) use this property to introduce a relaxation by optimising this PSSM in continuous space using the gradients of *l*. Concretely, the relaxed problem is

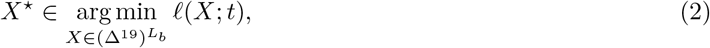

where *l* is extended to PSSM inputs so that it agrees with the discrete loss when every column of *X* is one-hot. This has since become the standard approach to solving eq. (1) in the field (Candido et al., 2026; Ovchinnikov, 2022; Pacesa et al., 2025). It is important to realise that *X*^*⋆*^ is not itself a protein sequence and so must be discretised to obtain 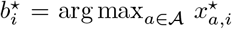, writing 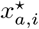 for the entry of *X*^*⋆*^ at amino acid *a* and position *i*. Additionally, we must now deal with a constrained optimisation problem, since *X* ranges over a product of simplices and each column must remain on the simplex.

In general the relaxed problem in eq. (2) is highly non-convex, so the design returned by a run of an optimiser depends on its random initialisation. We rank completed designs by a score 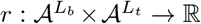, higher values being better, which is evaluated once per design and so, unlike *l*, need not be differentiable (section 4). A single run therefore produces a random design *b* whose score has survival function *G*(*u*) = Pr (*r*(*b*; *t*) ≥*u*). A campaign performs *n* independent runs but tests only the *m* highest-ranked designs, with *m* a small fraction of *n*, so performance depends on the upper tail of *G*, not its mean.

### 2.3 Loss

The hallucination-based binder design framework is extremely flexible; binders can be created for many different contexts by adding terms to the loss function. For example, binders can be designed for multiple targets simultaneously (Xu et al., 2026) or for specificity against off-targets (Brasas et al., 2026). Generally, a key component of the loss involves passing the candidate sequence, in complex with the target, through a structure prediction model, and using the model prediction to compute quantities related to the likelihood of the sequence being a binder to the target. Additional terms, such as perplexities from protein language models (Candido et al., 2026) or inverse folding model sequence recovery (Boyd et al., 2025; Dauparas et al., 2022), can also be added to regularise the sequence, and prevent adversarial solutions that show high structural confidence but are unlikely to work in practice due to poor solubility or expression.

In this work, we follow the loss introduced by Candido et al. (2026), which uses the ESM family of models: both the structure prediction model ESMFold2 and the protein language model ESMC. ESMFold2’s recurrent trunk applies successive transformer blocks to the ESMC embeddings of binder and target, producing a representation for a diffusion sampling head (final structure), a confidence head, and a distogram head. The structure losses read only the distogram *D*, which is a distribution over binned pairwise distances for every residue pair. The loss consists of four terms combined as a weighted sum (weights in appendix B):

- **Intra-chain** *l*_intra_: a contact-restricted cross entropy that favours confident short distances within the binder.
- **Cross-chain** *l*_inter_: the same contact loss, averaged over each target residue’s best binder contact, which encourages confident interface formation.
- **Globularity** *l*_glob_: a penalty on the predicted binder radius of gyration exceeding an empirical packing radius, which favours compact folds.
- **ESMC pseudo-log-perplexity** *l*_LM_: the mean cross entropy at randomly masked binder positions under ESMC, which regularises the design towards the language model’s prior.

### 2.4 Logit gradient descent

Softmax reparameterisation is a popular way to enforce the simplex constraints in hallucination-based binder design and is used by methods such as BindCraft (Pacesa et al., 2025). We note that some pipelines do not enforce the simplex constraint throughout: BindCraft, following AfDesign (Ovchinnikov, 2022), begins with a phase in which the model is fed a weighted sum of the raw logits and their softmax, which can place entries outside [0, 1], before later phases operate purely on the softmaxed representation, presumably to avoid the off-distribution nature of negative or greater-than-one amino-acid identities. We call the gradient-descent variant used by Candido et al. (2026) “Logit GD”, and it is the baseline against which we compare PGD. It introduces an unconstrained matrix of logits 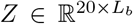 and defines the column-wise softmax map 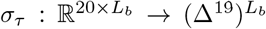, which sends the *i*-th column *z*_*i*_ of *Z* to softmax(*z*_*i*_*/τ*), where *τ >* 0 is a temperature. For any *Z*, every column of *σ*_*τ*_ (*Z*) is non-negative and sums to one, so *σ*_*τ*_ (*Z*) is feasible for eq. (2); the constrained problem thus becomes an *unconstrained* one over the logits, which can be solved using a standard gradient-based optimiser:

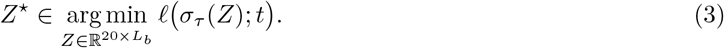

Candido et al. (2026) add several refinements to this basic method. Both the temperature *τ* ^(*k*)^ and the learning rate *α*^(*k*)^ are jointly annealed according to step *k* with a cosine schedule. Lowering the temperature causes the PSSM to sharpen and become closer to one-hot over the course of optimisation, so that by the final step only a single amino acid at each position has significant probability mass. This encourages minimal change in loss when discretising the PSSM and pushes the binder representation towards the one-hot representation that matches the structure prediction model’s training distribution. The learning rate must be jointly modified to maintain the same effective step size in the softmax-input space. Additionally, Candido et al. (2026) propose normalisation of the gradient of the loss, using separate normalisers for the structure and sequence terms in the loss, to ensure they always contribute to the total gradient at a fixed ratio. Pseudocode for our Logit GD implementation is given in appendix A.

The softmax reparameterisation method suffers from a pathology caused by the transformation. Figure 2 illustrates the geometry of the softmax transformation. Consider the gradient of the loss with respect to the logit for amino acid *a* at position *i*:

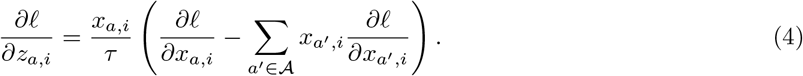

**Figure 2:**
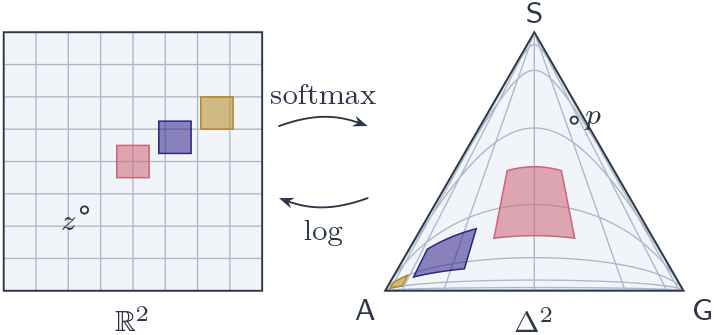
Softmax maps Euclidean logits onto the simplex. For |*A*| = 3, softmax is a bijection from the plane 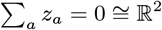 onto the interior of Δ^2^. Congruent squares (shaded) shrink towards the boundary: the map compresses space there, leaving gradients with respect to *z* poorly conditioned.

Here *x*_*a,i*_ is the probability assigned to amino acid *a* at position *i*, and *∂l/∂x*_*a,i*_ is the corresponding probability-space gradient. At fixed *τ >* 0, provided the probability-space gradients remain bounded, *∂l/∂z*_*a,i*_ tends to zero as *x*_*a,i*_ tends to zero. Thus an amino acid assigned very little probability receives little direct signal in logit space, even if increasing its probability would reduce the loss. As the whole column approaches a vertex, all of its logit-gradient components tend to zero. Normalising each loss gradient across the logit matrix can restore its overall norm but cannot undo unequal scaling across amino acids and positions.

## 3 Projected gradient descent

Whereas the softmax reparameterisation removes the constraints by optimising in an unconstrained logit space, projected gradient descent (PGD) keeps the optimisation in probability space and enforces feasibility with a projection step onto the probability simplex immediately following each update. The mosaic software package (Boyd et al., 2025) provides the first public implementation of simplex PGD for binder design, but no controlled comparison against alternative optimisers has been reported.

An ordinary gradient step ignores the constraints and generally leaves the feasible set, giving columns with negative or unnormalised entries. PGD simply follows each step with the Euclidean projection 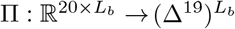 (Calamai & Moré, 1987; *Rosen, 1960), which acts independently on each column, replacing a column y* with the closest valid probability vector, 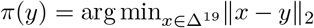. If the column already satisfies the constraints, it is unchanged. Each update is given by

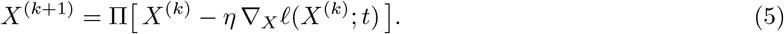

The projection can be computed exactly and efficiently, even in high dimensions, using the method of Duchi et al. (2008) and is cheap relative to a single evaluation of ∇_*X*_ *l*. Unlike the softmax map, which assigns every amino acid a strictly positive probability, the Euclidean projection can set probabilities exactly to zero. Applied to a column, the algorithm of Duchi et al. (2008) subtracts a common threshold *θ* from every entry and clamps the result at zero, *π*(*y*)_*a*_ = max{*y*_*a*_ − *θ*, 0}. The threshold is determined by finding the *θ* such that 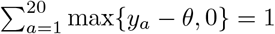. Pseudocode for our PGD implementation is given in appendix A.

We have described an alternative to the softmax reparameterisation method used by Candido et al. (2026), but the question remains: why should we expect this optimiser to perform better for the problem setup described in section 2.1? In binder design, the preference for an amino acid depends on the evolving sequence context. By updating probabilities directly, PGD avoids the softmax-induced attenuation of gradient information for low-probability amino acids (eq. (4)). Unlike Logit GD, the projection step means PGD can produce exact zeros, allowing optimisation to concentrate on a subset of amino acids while also ensuring that zeroed amino acids can be reintroduced if subsequent gradients favour them. Classical active-set identification results show that increasing sparsity throughout optimisation is not guaranteed (Calamai & Moré, 1987). This may allow for continued exploration compared with temperature-annealed Logit GD, which encourages probabilities to approach one-hot vectors by the end of optimisation.

## 4 Experimental setup

In this section we describe the experimental setup we use to evaluate the PGD optimiser. All experiments are in silico, and we do not carry out any in vitro testing of binding affinities. As motivated in section 2.1, we generate a large pool of designs and evaluate those in the upper tail. Unless otherwise specified, we run 1,024 designs per target/optimiser combination.

### Targets

We design binders for ten targets: human PD-L1 (UniProt Q9NZQ7), IL-7R*α* (P16871), TREM2 (Q9NZC2), RBX1 (P62877), TrkA/NTRK1 (P04629), VEGF-A (P15692), and TNF-*α* (P01375); Epstein– Barr virus BHRF1 (P03182); *E. coli* maltose-binding protein (MBP; P0AEX9); and the de novo designed beta barrel BBF-14 (PDB 9HAG). These targets have also featured in recent binder-design campaigns (Claude Science & Shanehsazzadeh, 2026; Pacesa et al., 2025; Watson et al., 2023). Exact target constructs are specified in appendix D.

### Design pipeline

Our design pipeline closely follows the ESMFold2 binder design pipeline presented in Candido et al. (2026). Consistent with that pipeline, we use the loss described in section 2.3. Both optimisers run for 150 steps, with cysteine excluded from the design alphabet. At each iteration, we randomly select one of two model variants trained on different datasets. During the final 20 optimisation steps, we also evaluate the interface confidence score ipTM (Evans et al., 2021) and retain the iterate with the highest ipTM. To make our final results more comparable to those of Claude Science & Shanehsazzadeh (2026), who design binders for the same targets and report experimental binding data for them, we depart slightly from the ESMFold2 pipeline by applying SolubleMPNN inverse folding to each optimised design, so that the sequences we rank are inverse-folded designs as in their campaign. We convert the selected iterate to a discrete sequence and predict the resulting sequence’s structure with both design models. The prediction with the higher ipTM supplies the reference backbone for the structural comparison score scDockQ (Basu & Wallner, 2016) and inverse folding using SolubleMPNN (Dauparas et al., 2022; Goverde et al., 2024). The resulting sequence is then ranked. We do not use filters to eliminate designs (Pacesa et al., 2025). A small parameter scan confirmed our choice of a PGD step-size scale of 0.1 and zero momentum (appendix C). In the PGD arm we retain the operations proposed by Candido et al. (2026), such as independent gradient scaling for the structure and sequence loss components, and we match the two arms as well as possible so that differences in design quality can be attributed to the optimiser. Further implementation details are given in appendix B.

### Inverse folding

Inverse folding takes the 3D structure of a protein backbone and predicts a sequence likely to fold into it. It is widely used in de novo sequence design (Wicky et al., 2022) and binder design (Cho et al., 2025a;b; Claude Science & Shanehsazzadeh, 2026; Frank et al., 2024; Jendrusch et al., 2025; Pacesa et al., 2025; Rustamov & Baev, 2025; Stark et al., 2025; Watson et al., 2023). Several studies attribute improved experimental success rates to this step, and report poor solubility in designs where it is omitted (Bennett et al., 2023; Frank et al., 2024; Wicky et al., 2022). We sample one sequence per optimised backbone using SolubleMPNN (Goverde et al., 2024) at temperature 0.1, keeping target sequences fixed and excluding cysteine. This differs from the ESMFold2 design methodology in Candido et al. (2026), which applies no inverse folding and instead filters designs on their isoelectric point.

### Ranking and selection

We primarily rank designs using the ipSAE score based on structure predictions from an ensemble of three models: ESMFold2, ESMFold2Fast, and ProtenixV2 (Candido et al., 2026; Zhang et al., 2026). Model settings are given in appendix B. For each model we run five separate predictions with different seeds. We compute the minimum of the target–binder and binder–target ipSAEs for each seed, then the maximum ipSAE among seeds for each model, and finally the mean across models. We also compute the self-consistency DockQ (scDockQ) score between the structure predicted by the design model and that predicted by the ranking model following SolubleMPNN inverse folding, using the highest-ipSAE seed for each model and averaging across models. This setup follows Claude Science & Shanehsazzadeh (2026), who provide evidence that it is predictive of wet-lab binding. For more detail, see appendix G.

### Metrics

We are interested in the ability of optimisers to produce high-quality designs in the upper tail of the score distribution (section 2.1). For each target and optimiser, we report the mean ensemble ipSAE among the top 10% of designs ranked by ensemble ipSAE, and the mean scDockQ among the top 10% ranked by scDockQ. Each metric therefore uses its own selected pool of 103 designs out of 1,024, rounding up. We also count in silico “hits”, defined as designs with ensemble ipSAE above 0.65 among all 1,024 designs. We use this fixed cutoff only as a reference for comparing optimisers. The survival curves in fig. 3 show the number of designs exceeding every ipSAE threshold, so the comparison does not depend on our chosen threshold.

**Figure 3:**
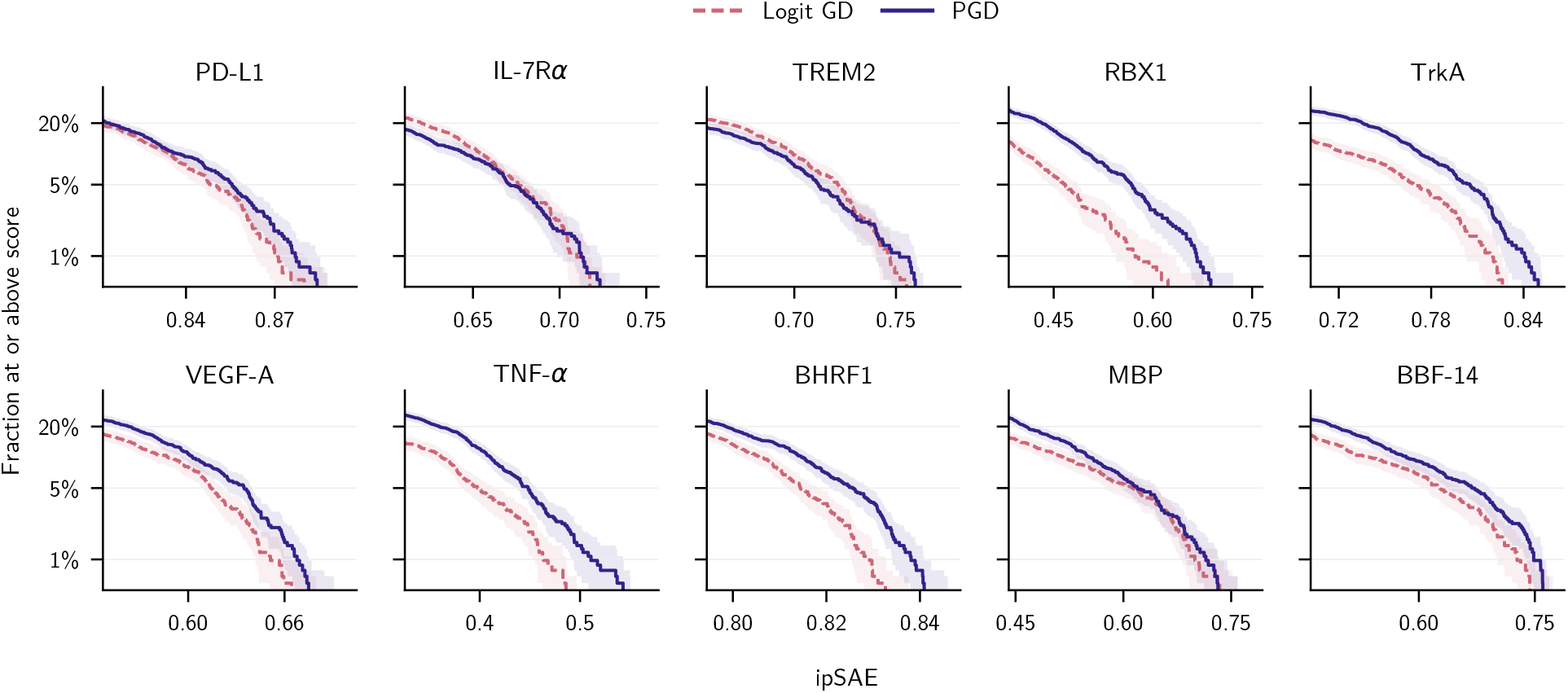
PGD increases the number of high-ipSAE binders. Survival curves for the upper tail of ensemble ipSAE for each target. Dashed is Logit GD; solid is PGD. Shading shows pointwise 95% nonparametric-bootstrap intervals, computed from their exact binomial marginals. The vertical axis uses a logarithmic scale.

### Statistics

We use paired bootstrap comparisons, matching designs across optimiser arms by their recorded design keys. Paired runs share their initial logits and per-step stochastic draws. We use bootstrapping to obtain uncertainty estimates for key metrics by resampling paired designs and reselecting the top 10% separately for each metric and optimiser 5,000 times. We report 95% confidence intervals. We report significant improvement on a per target basis for a metric when the confidence intervals on the paired difference do not contain 0.

### Compute

All experiments were conducted using NVIDIA GH200 GPUs on Isambard-AI (McIntosh-Smith et al., 2024). The primary run of both optimisation methods across all ten targets used a total of 1,377.29 GPU-hours of compute. For a per-target breakdown, see appendix E.

### Code and data availability

The design, ranking, and analysis code is available at github.com/ magnusross/pgd-hallucination. We release sequences, predicted structures, optimisation trajectories, scores, settings, and compute timings from all experiments at doi.org/10.5522/04/34493751 (Ross, 2026), together with analysis tables and figures (approximately 39 GiB compressed). The archive includes a README describing its contents; the code repository provides instructions for reproducing the analyses.

## 5 Results

In this section, we present experiments illustrating the favourable properties of the PGD optimiser in terms of efficiency, design quality, and design diversity.

### 5.1 Design quality

Figure 3 shows survival curves for the ensemble ipSAE values for the upper tail of a pool of 1,024 designs for each target. The upper tail is most relevant for real campaigns (section 2.1); full curves are shown in fig. 8. Across six of the ten targets (RBX1, TrkA, VEGF-A, TNF-*α*, BHRF1 and BBF-14), PGD provides significantly higher top-10% ensemble ipSAE, with the improvement for RBX1, TrkA and TNF-*α* being particularly large. On the remaining targets, the difference in top-10% ensemble ipSAE is not significant. Table 1 shows ensemble top-10% ipSAEs and scDockQs, as well as the number of in silico “hits” (section 4). PGD results in significantly better scores for both ipSAE and scDockQ for five of ten targets. PGD is never significantly worse than Logit GD on either scDockQ or ipSAE among the top 10% of designs. PGD provides more hits than Logit GD on five of ten targets; the methods are tied on two targets, and Logit GD provides more on the remaining three (IL-7R*α*, TREM2 and BHRF1). The improvement in hit counts for PGD is particularly large on some “harder” targets, where ipSAEs are lower; in particular PGD provides over 5× the number of hits on RBX1 (17 versus 3), and nearly 2× on VEGF-A (23 versus 12); the greatest proportional improvement of Logit GD is only 1.24×, on IL-7R*α* (117 versus 94). These ratios are descriptive and, for RBX1, depend on a small number of hits; neither method produces hits on TNF-*α*. Further results, including median statistics, can be found in appendix H.

**Table 1:** Top-10% mean ipSAE and scDockQ (103 selected designs per method and target), and numbers of designs with ipSAE *>* 0.65 among all 1,024 designs. Bold identifies the higher method only when its 95% paired-bootstrap interval excludes zero. Hit counts have no significance marking.

| Measure | Method | PD-L1 | IL-7Rα | TREM2 | RBX1 | TrkA | VEGF-A | TNF-α | BHRF1 | MBP | BBF-14 |
| --- | --- | --- | --- | --- | --- | --- | --- | --- | --- | --- | --- |
| Top-10% ipSAE | Logit GD | 0.852 | 0.681 | 0.723 | 0.484 | 0.774 | 0.620 | 0.410 | 0.817 | 0.615 | 0.637 |
|  | PGD | 0.858 | 0.678 | 0.718 | **0.578** | **0.807** | **0.636** | **0.456** | **0.826** | 0.632 | **0.672** |
| Top-10% scDockQ | Logit GD | 0.910 | 0.842 | 0.858 | 0.487 | 0.859 | 0.877 | 0.618 | 0.907 | 0.821 | 0.751 |
|  | PGD | 0.914 | 0.828 | 0.849 | **0.522** | **0.882** | **0.894** | **0.669** | 0.909 | **0.841** | **0.783** |
| ipSAE $> 0.65$ | Logit GD | 633 | 117 | 239 | 3 | 210 | 12 | 0 | 797 | 35 | 41 |
|  | PGD | 658 | 94 | 198 | 17 | 360 | 23 | 0 | 771 | 35 | 61 |

### 5.2 Structural diversity

The diversity of candidates in the selected pool is also an important concern in any design campaign; even if scores are high, binders with similar structures may share experimental failure modes. Binder chains from the ESMFold2 ranking model predictions with the best ipSAE from the five seeds were clustered using Foldseek (van Kempen et al., 2024) at TM-score 0.6, using alignment-length normalisation and setting the sequence-identity and coverage cutoffs to zero. Figure 4 shows the number of clusters for each target. These counts describe similarity to cluster representatives at this threshold, rather than distinct biological folds. PGD produces sequences that are much more structurally diverse than those produced by Logit GD. For every target, PGD has higher cluster counts, often more than twice as many. We hypothesise that this improvement comes from the difference in optimiser dynamics, to which we turn in the next section.

**Figure 4:**
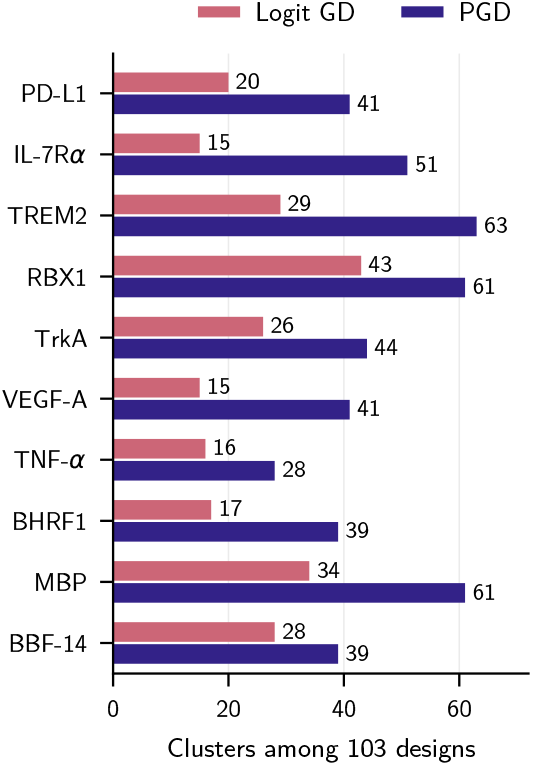
PGD designs are more structurally diverse. Foldseek cluster counts among the top-10% designs by ipSAE.

### 5.3 Optimisation dynamics

In fig. 5 we plot a number of quantities related to the optimisation dynamics of each method, which suggest an explanation for the superior performance of PGD, and are consistent with the theoretical benefits described in section 3. Panel (a) shows that PGD achieves a greater mean decrease in objective relative to step 1 over the course of optimisation, and is able to descend much more steeply than Logit GD early in optimisation. In (b) we plot the mean number of active amino-acid choices (probability *>* 10^*−*3^) at each position over optimisation. The temperature annealing used in Logit GD pushes the iterate towards a vertex so that the probability mass is focussed on a single amino acid by the end of optimisation, whereas with PGD, sparsity does increase, but an active set of around 10 amino acids per position is maintained in later optimisation steps. Panel (c) illustrates the pathology discussed in section 2.4; once we stray near an edge with Logit GD, we stay there; amino-acid choices whose probability falls below 10^*−*3^ essentially never become active again. PGD avoids this issue, which may allow it to explore more of sequence space. In (d) we can see that although PGD retains soft iterates in the final selection window, its argmax sequences achieve lower mean objective values than those from Logit GD. For both methods, the argmax sequences have a worse mean objective than the soft iterates but this should not be attributed to the cost of discretisation: the soft iterate is chosen as the best of several stochastic evaluations in the tail of the trajectory, so its recorded measurement is favourably selected, whereas the discrete sequence’s measurement is a single unbiased draw. Some of the gap is therefore expected even if discretisation has no expected cost.

**Figure 5:**
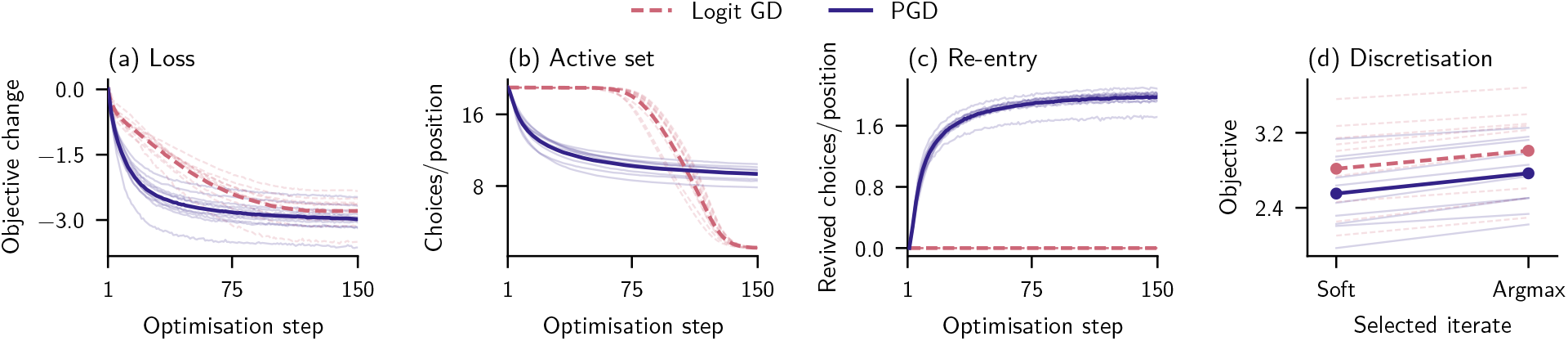
PGD maintains flexibility during optimisation whilst reaching better objectives. In all plots, translucent lines show the mean across designs for different targets; prominent lines are the mean across targets. (a) Objective change relative to step 1. (b) Active amino-acid choices per position (probability *>* 10^*−*3^). (c) Choices per position that move from probability ≤ 10^*−*3^ to *>* 10^*−*3^ on the next step. (d) Soft and argmax objectives at the tail-selected iterate.

### 5.4 Sensitivity to campaign details

We chose the top-10% (103 designs) as a simulated selection pool; assays can often be batched to around this number. In fig. 6 we show that PGD’s higher mean ipSAE across targets is not sensitive to the specific choices of selection fraction and pool size tested. Panel (a) shows mean ipSAE for the selected top fraction, for selection fractions from 1% to 20%, averaged across targets. PGD provides higher mean ipSAE across all tested selection fractions. Panel (b) shows how the expected ipSAE of the best design changes for each method as the pool size is increased from 32 to 1,024. We compute the exact expected maximum of a uniformly sampled subset from the full pool, without replacement, at each size, and then compute the mean across targets. The expected ipSAE of the best design, averaged across targets, is higher for PGD at every tested pool size.

**Figure 6:**
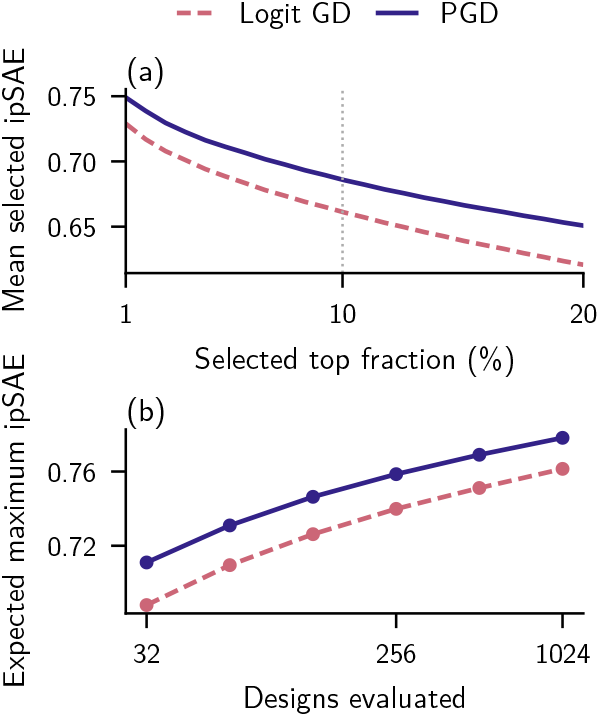
PGD provides better designs for different campaign requirements. Mean ipSAE across targets for (a) the selected top fraction of each 1,024-design pool and (b) the expected maximum of a uniformly sampled subset, without replacement. The dotted line marks 10% selection.

### 5.5 Better designs or more designs?

The curves in fig. 5 (a) indicate that PGD is able to achieve good objective values rapidly. This suggests that more high-quality designs might be obtained for the same amount of compute by running more designs for fewer optimisation steps. We compare the standard 150-step policy, which ranks all 1,024 candidates per target^1^, with a 75-step PGD policy that generates 2,048 candidates and only applies the full ranking to the top half selected by one-seed ESMFold2Fast ipSAE after SolubleMPNN redesign of all candidates. We also halve the number of tail selection steps, so compute is similar; screening and compute-accounting details are given in appendix F. We measure computational yield as the number of ranked designs exceeding an ipSAE threshold divided by recorded design and ranking GPU-hours, including redesign and screening of all candidates. Across the ten targets with available rankings, pooled yield at ipSAE *>* 0.65 rises from 3.29 hits per GPU-hour for 150-step PGD to 4.24 for screened 75-step PGD, a 29% increase; 150-step Logit GD yields 3.08 (fig. 7). The shorter policy improves yield over both 150-step policies on eight of ten targets, with gains of 3–57% relative to the better of the two; only RBX1 declines, by 24% (fig. 7). Furthermore, 75-step PGD gives a higher yield than Logit GD on nine of ten targets, with a tie on TNF-*α*, where neither policy produces hits.

**Figure 7:**
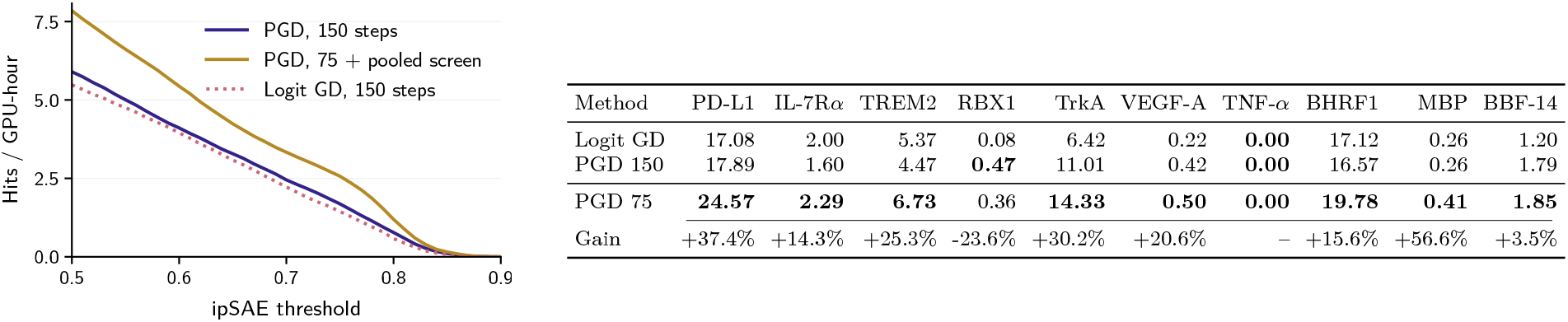
More short PGD runs increase computational yield. Hits per GPU-hour pooled across ten targets with available rankings (left) and per target at ipSAE *>* 0.65 (right). The final row gives PGD 75’s percentage gain over the better of the two 150-step methods. Bold values mark the highest yield per target, without significance testing.

## 6 Limitations and future work

The primary limitation of this work is that no wet-lab validation of binders was conducted. Although ipSAE has been shown in numerous analyses to be a reasonably good predictor of real binding (Claude Science & Shanehsazzadeh, 2026; Overath et al., 2025), there remains the possibility that some aspect of the PGD designs makes them unsuitable. The analysis presented here is limited to a single pipeline (Candido et al., 2026), chosen for its strong performance in binder generation, and its recency. The nature of the problem, i.e. that success hinges on upper-tail performance, means that we must generate many designs to get a reliable statistical effect, requiring a large amount of compute. Although early experimentation provided evidence for generalisation across pipelines, we chose to prioritise a reliable result on a single pipeline over partial results for multiple pipelines. We believe the mechanism presented in fig. 5 is likely to generalise. The same compute requirement also meant that we were not able to conduct extensive ablations across algorithm hyperparameters or experimental setups.

The results measure predicted interface confidence rather than binding. Furthermore, ESMFold2 and ESM-Fold2Fast, two of the three structure predictors that scored the designs, share the design model family. The ipSAE *>* 0.65 threshold that we chose for potential binder hits is not calibrated per target and is a computational proxy for experimental success rather than a measured hit rate. Additionally, we only compared PGD with Logit GD, and did not compare with other constrained optimisation methods such as mirror descent and sparsemax.

We made a choice to pass each design through SolubleMPNN inverse folding before it is ranked. We adopted this choice to make our designs comparable to the experimentally validated campaign of Claude Science & Shanehsazzadeh (2026) and because inverse folding is widely reported to improve solubility (section 4). One drawback is that the final sequence that we score is not the sequence directly produced by the optimiser, as such, any advantage of the optimiser must be transmitted through the inverse folding step, which may attenuate the differences we measure. Nevertheless, we are reassured to see that the preference for PGD is visible prior to inverse folding: we note that the argmax-sequence objective at the selected iterate is lower for PGD than for Logit GD (fig. 5d).

We believe that the optimisation part of hallucination-based binder design has been neglected to date, and there are potentially big efficiency improvements to be had by leaning on existing literature. We have provided evidence for one such algorithm here, but there is likely much further to go.

## 7 Conclusion

In this work, we have presented projected gradient descent as a performant, efficient alternative algorithm for hallucination-based binder design. We have shown that it is able to generate higher-quality designs than the popular approach of gradient descent on softmax logits. Across ten targets with 1,024 designs per target and optimiser, PGD gave significantly higher top-10% ensemble ipSAE on six of ten targets, was significantly lower on none, and produced more structurally diverse designs on all ten. We provide evidence for the mechanism behind this performance, which indicates that projected gradient descent is able to more extensively explore sequence space. We find that for most targets, running more designs whilst using less compute on each is more computationally efficient.

## Acknowledgements

The authors acknowledge the use of resources provided by the Isambard-AI National AI Research Resource (AIRR). Isambard-AI is operated by the University of Bristol and is funded by the UK Government’s Department for Business, Innovation, Science and Trade (DBIST) via UK Research and Innovation; and the Science and Technology Facilities Council [ST/AIRR/I-A-I/1023]. M.R. is supported by the EPSRC Digital Health Hub for Antimicrobial Resistance (EP/X031276/1). D.M. is supported by funding from the MRC (MR/W006774/1). J.W. is supported by ARIA’s Activation Partnership with Pillar VC. We thank Nick Boyd and Sam Guns for informative advice.

## A Optimisation algorithms

Algorithm 1 gives simplified pseudocode for the two optimisers. Both use the same initial logits *Z*^(0)^, exclude cysteine, and share the random model choices and stochastic draws at each step. Here *l*_*S*_ is the weighted sum of the three structure losses and *l*_LM_ is the unweighted ESMC loss (section 2.3). We use *T* = 150 steps, language-model weight *λ* = 0.15, and PGD step size 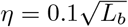.

### Algorithm 1

Logit GD and PGD for binder design. Blue lines highlight where the optimisers differ.

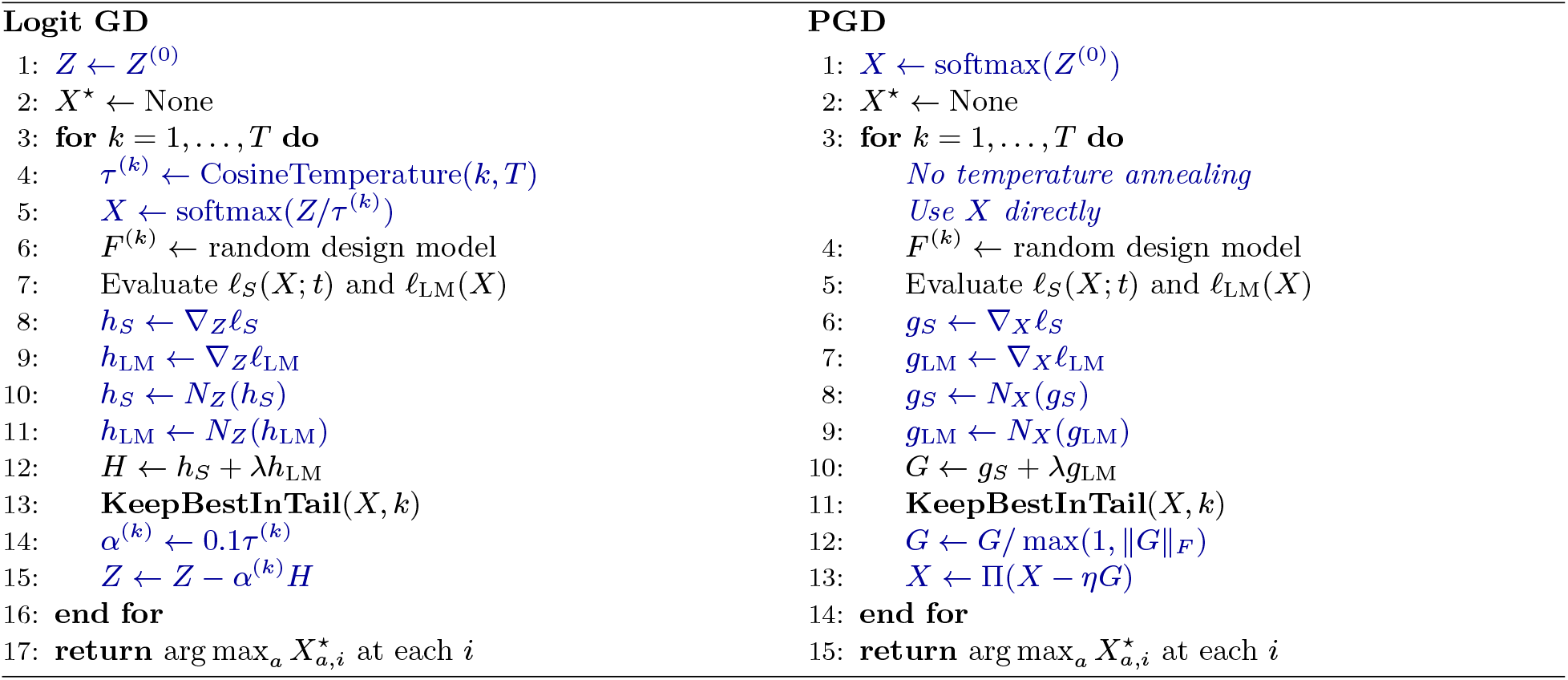

### Normalisation and updates

*N*_*Z*_ normalises each logit-gradient matrix to Frobenius norm *L*_effective_, where *L*_effective_ counts positions with nonzero gradient. *N*_*X*_ first subtracts the amino-acid mean at each position, then normalises the probability-gradient matrix to unit Frobenius norm. Thus Logit GD normalises *after differentiating through softmax*, whereas PGD normalises *in probability space*, clips the combined direction, and projects. The Euclidean projection Π of eq. (5) acts independently at each position and keeps cysteine at zero. The cosine temperature schedule is specified in appendix B.

### Shared tail selection

**KeepBestInTail** evaluates ipTM on the current soft iterate using *F* ^(*k*)^ during the final 20 steps, and updates *X*^*⋆*^ whenever this score exceeds the best previously observed in that window. Discretisation of the soft sequence only happens after all optimisation steps are complete and is applied only to the best evaluated iterate in the tail. For 75-step PGD, the selection window is the final ten steps. Subsequent reference prediction, inverse folding, and ranking follow section 4.

## B Implementation details

### Optimisers

Both arms use paired initial logits drawn from *N*(0, 0.01^2^) for allowed amino acids; PGD starts from their softmax probabilities. Logit GD uses the cosine temperature schedule *τ* ^(*k*)^ = 0.01 +0.99[1 + cos(*πk/*150)]*/*2 for *k* = 1, …, 150. Gradient normalisation, step sizes, and tail selection are compared in appendix A.

### Objective and design inference

We use the miniprotein objective, weights, and loss constants in Algorithms 11–14 of Candido et al. (2026). Design uses the ESMFold2-Experimental-Fast and ESMFold2-Experimental-Fast-Cutoff2025 checkpoints, which are the two variants sampled at each optimisation step, with one recycle, language-model dropout 0.5, and no target MSA. Tail ipTM evaluation and final reference predictions use 50 diffusion sampling steps and one sample per prediction.

### Inverse folding and scoring

SolubleMPNN uses checkpoint soluble_v_48_020. We compute ipSAE using Mosaic’s IPSAE_min implementation with a PAE cutoff of 10 Å, grouping all target protein chains together and excluding ligand tokens. DockQ uses positional residue correspondence with target chains merged, subject to the TREM2 residue restriction described in appendix D. For homomeric targets, we maximise scDockQ over permutations of identical target chains.

References with no interface contacts receive scDockQ zero.

### Ranking model settings

We use 3 recycles and 50 diffusion sampling steps for ESMFold2 and ESM-Fold2Fast, and 10 recycles and 200 diffusion sampling steps for ProtenixV2, with one sample per seed. Target MSAs are used by ESMFold2 and ProtenixV2 but not ESMFold2Fast.

## C PGD parameter scan

We checked the PGD step-size scale and momentum on seven targets: PD-L1, IL-7R*α*, TREM2, RBX1, TrkA, BHRF1, and MBP. The hyperparameter sweep tested 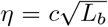 with *c* ∈ {0.05, 0.075, 0.1, 0.15} and momentum *m* ∈ {0, 0.25}, using 150 optimisation steps and 128 designs for each target and setting, with the same ranking pipeline. We evaluated using top-10% ensemble ipSAE and scDockQ. Nearby step-size scales gave similar performance. Momentum provided no consistent benefit across targets. This supported retaining *c* = 0.1 and *m* = 0 for the main experiments.

## D Target constructs

Table 2 lists the target inputs used for design and ranking. For all targets, we design binders with a fixed length of 80 residues. VEGF-A and TNF-*α* are modelled as a homodimer and homotrimer, respectively. RBX1 includes three Zn ions in structure prediction; these are excluded from protein losses, confidence metrics, DockQ, and SolubleMPNN input. Our TREM2 construct contains the folded Ig-like domain and a flexible extracellular stalk (Szykowska et al., 2021). TREM2 uses all 156 residues for prediction and ipSAE; for scDockQ (Basu & Wallner, 2016), we crop both reference and predicted target structures to construct residues 1–112 (UniProt residues 19–130), excluding the stalk from both contact and RMSD calculations.

**Table 2:** Target constructs. Lengths count residues per target protein chain, excluding the 80-residue binder and ligand tokens. Accessions refer to UniProt, except BBF-14 (PDB).

| Target | Accession | Protein chains | Residues/chain | Ligands |
| --- | --- | --- | --- | --- |
| PD-L1 | Q9NZQ7 | 1 | 115 | None |
| IL-7Rα | P16871 | 1 | 193 | None |
| TREM2 | Q9NZC2 | 1 | 156 | None |
| RBX1 | P62877 | 1 | 108 | 3 Zn ions |
| TrkA | P04629 | 1 | 101 | None |
| VEGF-A | P15692 | 2 | 94 | None |
| TNF-α | P01375 | 3 | 157 | None |
| BHRF1 | P03182 | 1 | 157 | None |
| MBP | P0AEX9 | 1 | 370 | None |
| BBF-14 | 9HAG (PDB) | 1 | 110 | None |

## E Compute breakdown

Table 3 reports allocated GPU-hours for the 320 completed cells that together contain 1,024 designs per target and optimiser (20,480 designs in total). Compute is measured as task wall time multiplied by allocated GPU count, including setup and CPU work within each task but excluding queue time, failed attempts, and other optimiser recipes. Ranking includes SolubleMPNN redesign and subsequent evaluation. Design used 943.61 GPU-hours and ranking used 433.69 GPU-hours; total compute was highest for TNF-*α* (415.73 GPU-hours) and lowest for TrkA (65.43 GPU-hours).

**Table 3:** Allocated GPU-hours per target, summed over both optimisers. Totals are calculated before rounding.

| Target | Design | Ranking | Total |
| --- | --- | --- | --- |
| PD-L1 | 49.14 | 24.70 | 73.84 |
| BBF-14 | 45.78 | 22.56 | 68.34 |
| BHRF1 | 63.34 | 29.76 | 93.10 |
| IL-7Rα | 80.85 | 36.25 | 117.11 |
| MBP | 186.89 | 83.73 | 270.62 |
| RBX1 | 46.78 | 27.28 | 74.06 |
| TNF-α | 293.32 | 122.41 | 415.73 |
| TREM2 | 60.23 | 28.55 | 88.77 |
| TrkA | 43.54 | 21.88 | 65.43 |
| VEGF-A | 73.74 | 36.55 | 110.29 |
| Total | 943.61 | 433.69 | 1,377.29 |

## F Short-run screening and compute

For each target, the 75-step policy redesigns all 2,048 candidates with SolubleMPNN, then screens them with one ESMFold2Fast seed, using 3 recycles and 50 diffusion steps. The 1,024 candidates with the highest ipSAE are then selected to receive full ensemble ranking. Due to compute budget constraints, TNF-*α* uses 896 designs per policy that go on to final ranking rather than 1,024. Compute estimates for both arms include all design, redesign, and screening costs. Table 4 reports the resulting compute budgets, including task setup and CPU work.

**Table 4:** Compute and hit counts for the matched allocation comparison. Hits have ensemble ipSAE *>* 0.65. Each policy contributes 1,024 ranked designs per target, except TNF-*α* (896).

| Policy | Hits | GPU-hours |
| --- | --- | --- |
| Logit GD, 150 steps | 2,087 | 676.65 |
| PGD, 150 steps | 2,217 | 674.65 |
| PGD, 75 steps + screening | 3,087 | 727.93 |

## G In silico scores and experimental binding

Bennett et al. (2023) was one of the first works to show that filtering de novo binder designs using AlphaFold2’s inter-chain Predicted Aligned Error (PAE) scores improves experimental hit rates by approximately a factor of ten. Metrics derived from the PAE matrix, such as ipTM (Evans et al., 2021) and later ipSAE (Dunbrack, 2025), have been repeatedly shown to be good predictors of experimental binding. Overath et al. (2025) found that ipSAE from AlphaFold3 was the single best predictive metric in retrospective analysis of 3,766 experimentally tested de novo candidate binders across 15 targets, outperforming ipAE and ipTM. In a reanalysis of 13 targets from that dataset, Claude Science & Shanehsazzadeh (2026) evaluated an ensemble of ESMFold2, ESMFold2Fast, and ProtenixV2, reporting a mean within-target average precision (macro-AP) of 0.66 for the model-ensembled ipSAE and 0.65 for the model-ensembled combination of ipSAE and scDockQ, against a background mean per-target hit rate of 0.15.

Following this retrospective evaluation, Claude Science & Shanehsazzadeh (2026) used the combined ipSAE and scDockQ score to select designs for prospective testing, obtaining interpretable results for 15 targets, including all ten studied here. The overall hit rate was 26.8% (354/1,320), with a median per-target hit rate of 22.2%. Even within these preselected pools, subsequent rescoring with the three-model ipSAE ensemble, standardised within target, distinguished hits from non-hits: average precision exceeded chance on 12 of 13 evaluable targets (sign test, *p* = 0.003), averaging 0.52 against a chance baseline of 0.31. These results support using the ensemble ipSAE as a proxy for increased binding probability, including among already high-ranking designs.

In a meta-analysis of experimentally tested de novo binders, the F1-maximising threshold for AlphaFold3 ipSAE was 0.61, with per-target thresholds mostly between 0.5 and 0.8 (Overath et al., 2025). This variation motivates reporting score distributions alongside hit counts at a fixed threshold, rather than treating a single cutoff as a target-independent predictor of experimental success.

## H Additional results

Figure 8 shows the full ensemble ipSAE survival curves for each target and optimiser. Table 5 reports median ensemble ipSAE and scDockQ across the full design pools.

**Figure 8:**
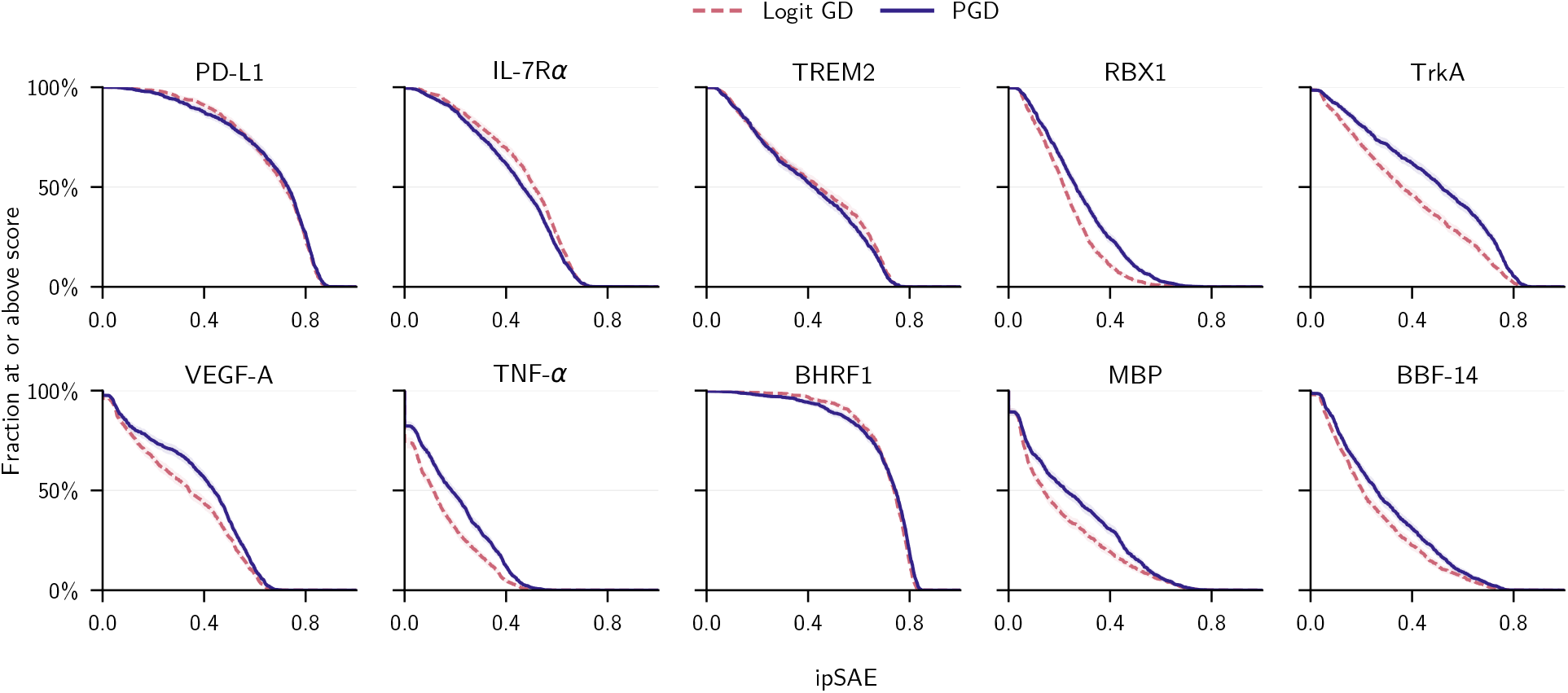
Full survival curves for ensemble ipSAE across all 1,024 designs per target and optimiser. Dashed is Logit GD; solid is PGD. Shading shows pointwise 95% nonparametric-bootstrap intervals, computed from their exact binomial marginals.

**Table 5:** Median ensemble ipSAE and scDockQ across all 1,024 designs per target and optimiser. These are descriptive statistics; no significance marking is applied.

| Measure | Method | PD-L1 | IL-7Rα | TREM2 | RBX1 | TrkA | VEGF-A | TNF-α | BHRF1 | MBP | BBF-14 |
| --- | --- | --- | --- | --- | --- | --- | --- | --- | --- | --- | --- |
| Median ipSAE | Logit GD | 0.715 | 0.510 | 0.442 | 0.219 | 0.358 | 0.341 | 0.115 | 0.738 | 0.139 | 0.205 |
|  | PGD | 0.730 | 0.465 | 0.421 | 0.267 | 0.514 | 0.438 | 0.185 | 0.745 | 0.225 | 0.259 |
| Median scDockQ | Logit GD | 0.755 | 0.521 | 0.504 | 0.241 | 0.432 | 0.534 | 0.243 | 0.766 | 0.336 | 0.370 |
|  | PGD | 0.770 | 0.474 | 0.471 | 0.281 | 0.587 | 0.663 | 0.386 | 0.756 | 0.454 | 0.440 |

## Footnotes

1 For compute reasons, TNF-α uses 896 ranked designs per method. Controls are subsampled without using scores to match this cohort.

